# permGWAS2: Enhanced and Accelerated Permutation-based Genome-Wide Association Studies

**DOI:** 10.1101/2023.11.28.569016

**Authors:** Maura John, Arthur Korte, Dominik G. Grimm

## Abstract

**Motivation:** Permutation-based significance thresholds have been shown to be a robust alternative to Bonferroni-based significance thresholds in genome-wide association studies (GWAS). However, the implementation of permutation-based thresholds is computationally demanding. The recently published method `permGWAS` introduced a batch-wise approach using 4D tensors to efficiently compute permutation-based GWAS. However, running multiple univariate tests in parallel leads to many repetitive computations and increased computational resources. More importantly, the previous version of `permGWAS` does not take into account the population structure when permuting the phenotype.

**Results:** We propose `permGWAS2`, an improved and accelerated version that uses a block matrix decomposition to optimize computations, thereby reducing redundant computations. It also introduces an alternative permutation strategy that takes into account the population structure during permutation. We show that this improved framework provides a more streamlined approach to performing permutation-based GWAS with a lower false discovery rate compared to the previous version and the traditional Bonferroni correction.

**Availability:** `permGWAS2` is open-source and publicly available on GitHub for download: https://github.com/grimmlab/permGWAS.

## 1 Introduction

Linear mixed models (LMMs) are a popular method for correcting for confounding factors, such as population structure and cryptic relatedness, when conducting genome-wide association studies (GWAS) (Lippert *et al*., 2011; Loh *et al*., 2015; Gumpinger *et al*., 2018). Permutation-based significance thresholds provide a robust alternative to Bonferroni thresholds that can limit false-positive associations in statistical hypothesis tests (Che *et al*., 2014; Nicod *et al*., 2016; John *et al*., 2022). One of the main challenges in using permutation-based approaches is the high computational cost, which was recently addressed in John *et al*. (2022). There we introduced `permGWAS`, a Python framework capable of computing efficient batch-wise LMMs using 3- and 4-dimensional tensors. This reformulation, combined with the *maxT* permutation method of Westfall and Young (1993), allowed us to compute permutation-based thresholds in a reasonable amount of time. However, running multiple univariate tests in parallel inevitably leads to increased memory usage. In addition, although `permGWAS` processes the SNPs in each batch simultaneously, it still performs each hypothesis test independently, repeating certain computations that could actually be saved and reused. In fact, when estimating the parameters of an LMM, certain calculations are independent of the marker of interest and can therefore be reused for each SNP. More importantly, `permGWAS` currently contains only a simple permutation strategy that does not take the population structure into account when permuting the phenotype.

Here we propose `permGWAS2`, an improved and accelerated version of `permGWAS` that takes advantage of a block matrix decomposition to reduce the number of redundant computations. In addition, our new framework now provides an alternative permutation approach that takes population structure into account.

## 2 Methods

In the following, we first review the basic framework of linear mixed models and how to estimate effect sizes and variance components, followed by two lemmas on how to efficiently compute permutation-based significance thresholds while taking into account the population structure during permutations.

### 2.1 Linear mixed models

For a vector of *n* phenotypic observations *y* ∈ ℝ^*n*^, we consider a linear mixed model (LMM) of the form

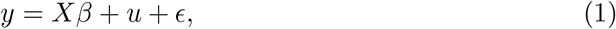

where *X* ∈ ℝ^*n×c*^ is a matrix of fixed effects including a column of ones for the overall mean, the covariates, and the SNP of interest; *β* ∈ ℝ^*c*^ denotes the effect sizes of the fixed effects; *u* ∈ ℝ are random effects with 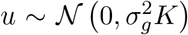, for the genetic variance component 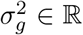 and the genetic similarity matrix *K* ∈ ℝ^*n×n*^; and *ϵ* ∈ ℝ^*n*^ is a vector of residual effects with 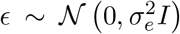 for residual variance component 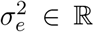 and identity matrix *I*∈ ℝ^*n×n*^. Then *y* also follows a Gaussian distribution with mean *Xβ* and covariance matrix 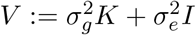.

#### Estimation of log likelihood

We estimate the effect sizes *β*, and the variance components 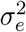and 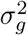 by maximizing the likelihood function

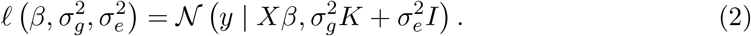

This is equivalent to finding the maximum of the log-likelihood function.

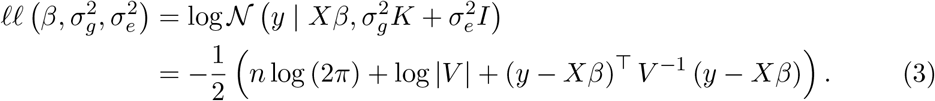

Let 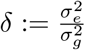 and *H* := *K* + *δI*. Then Equation 3 can be simplified to

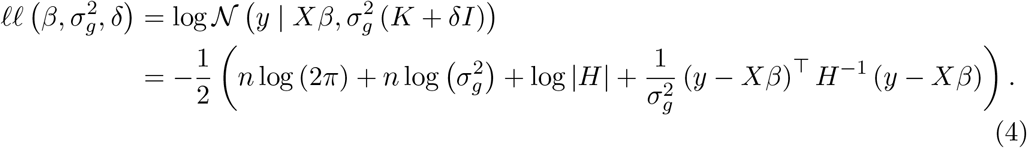

Computing the derivative with respect to *β* and setting it equal to zero yields the generalized least squares estimate:

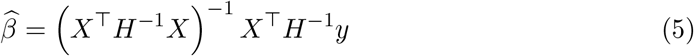

Substituting *β* in Equation 4 with 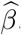, computing the derivative with respect to 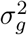 and setting it equal to zero yields

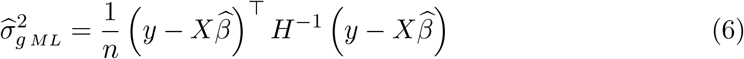

Finally we can substitute *β* and 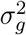 in Equation 4 with 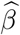 and 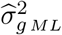, and the log likelihood becomes a function only dependent on *δ*:

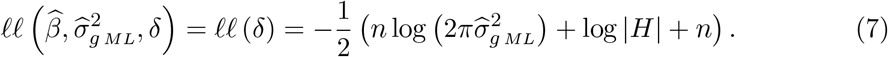

#### Efficient evaluation of log likelihood

Since the kinship matrix *K* is symmetric, we can compute the spectral decomposition *K* = *UDU*^⊤^, where *U* is an orthogonal matrix of eigenvectors of *K*, i.e., *UU*^⊤^ = *I*, and *D* is a diagonal matrix containing the corresponding eigenvalues *λ*_*i*_ for *i* ∈ {1, …, *n*}.

Then *H* = *U* (*D* + *δI*) *U*^⊤^ and *H*^−1^ = *U* (*D* + *δI*)^−1^ *U*^⊤^. Hence, we get

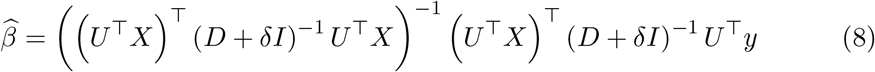

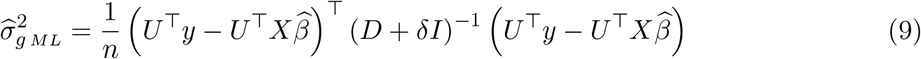

and by using the fact that |*U*| = 1, Equation 7 changes to

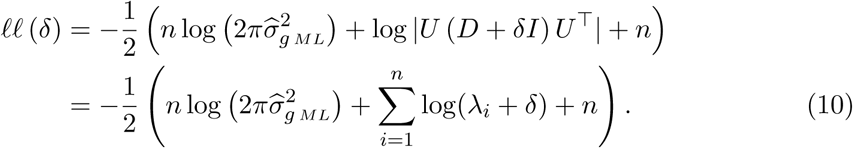

#### Restricted maximum likelihood

We can extend Equation 4 to get the restricted maximum likelihood (REML):

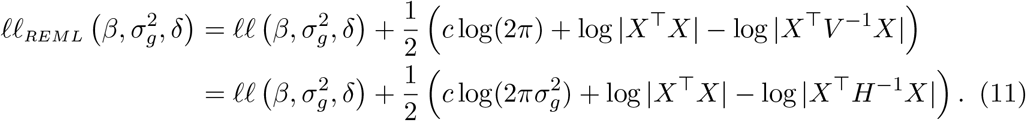

Then the formula for 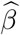 remains unchanged and the variance component estimate is given by

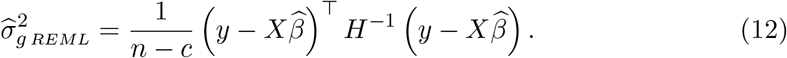

Similarly to equation 7 we can put 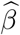 and 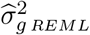 into Equation 11 and get

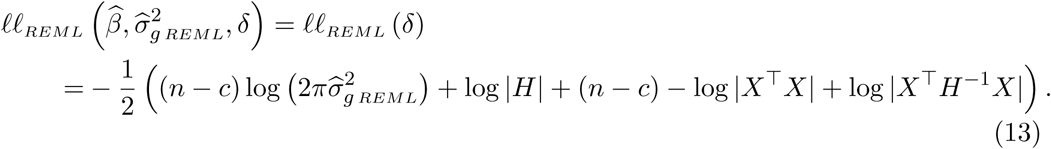

Using again the spectral decomposition of *K*, this can be efficiently reformulated to

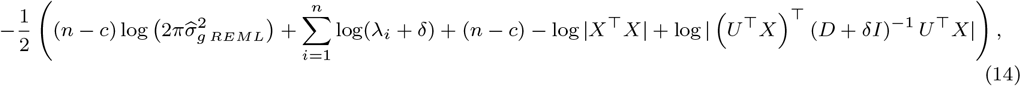

with

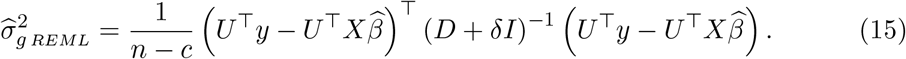

Hence, maximizing the log likelihood function 11 is equivalent to finding *δ* ∈ ℝ that maximizes Equation 14. In order to find the globally optimal *δ* we use an approach similar to Kang *et al*. (2008) and Lippert *et al*. (2011): First, we compute the function values with respect to Equation 14 of 100 equidistant values between 10^−5^ and 10^5^ on a logarithmic scale. Then, for each triplet of neighboring points where the function value in the middle is greater than at the boundaries, we apply Brent’s method to find the locally optimal *δ*. Finally, we take the optimal *δ* among all evaluated points. To speed up the computation, we estimate *δ* and 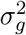 only once for a null model without any genetic markers, and reuse the estimates for the alternative models including the markers of interest as described in Kang *et al*. (2010). Once we have estimated the variance components, we use an F-test to test the null hypothesis that the marker of interest has no effect on a given phenotype. If the resulting p-value is below a predefined significance threshold, we reject the null hypothesis.

Because we test thousands to millions of markers at once in a typical GWAS, we need to correct for these multiple hypotheses to avoid thousands of false-positive associations. One way to do this is to empirically estimate the family-wise error rate, i.e., the probability of making at least one false positive, by computing a permutation-based significance threshold (John *et al*., 2022).

### 2.2 Permutation-based significance thresholds

To compute a permutation-based significance threshold for a given significance level *α*, we use the *maxT* method of Westfall and Young (1993). For this, we first permute the phenotype *y q*-times and compute the test statistics ^*k*^*t*_*i*_ for each permutation *k* ∈ {1, …, *q*} and marker *i* ∈ {1, …, *m*}. Then for each permutation, we take the maximal test statistic over all markers ^*k*^*t*_max_ := max_*i*∈{1,…,*m*}_^*k*^*t*_*i*_ which corresponds to the minimal p-value ^*k*^𝓅_min_. Finally, the adjusted threshold is given as the *α*-th percentile of the minimal p-values. This permutation-based threshold is able to control the family-wise error rate, as shown in John *et al*. (2022).

However, the permutation strategy presented above does not take into account the underlying population structure of the given phenotype. In fact, permuting the phenotype breaks not only the correlation between the phenotypic and genotypic values, but also the relatedness between the individuals. To preserve this relatedness, one can alternatively compute the covariance matrix anew for each permutation or, equivalently, permute the rows and columns of the covariance matrix using the same permutation as for the phenotype vector. By Lemma 1 this is equivalent to permuting the fixed effects matrix containing the covariates and the SNP of interest.

#### Lemma 1.

*Consider the LMM from the Equation 1, i*.*e*., *y* = *Xβ* +*u*+*ϵ with covariance matrix* 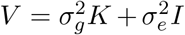. *Let τ* : {1, …, *n*} → {1, …, *n} be a permutation. Then permuting the entries of the phenotype vector y and the rows and columns of V with respect to τ is equivalent to permuting the rows of X with respect to the inverse permutation τ*^−1^.

*Proof*. Let *P*_*τ*_ ∈ ℝ^*n×n*^ denote the permutation matrix obtained by permuting the rows of the identity matrix *I* ∈ ℝ^*n×n*^ according to *τ*. Note that permutation matrices are orthogonal, i.e., 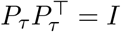, and that 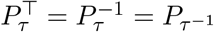 is the permutation matrix of the inverse permutation *τ*^−1^. Then multiplying a matrix *A* with *P*_*τ*_ from left permutes the rows of *A* and multiplying it from right with 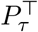 permutes the columns of *A* according to *τ*.

Now consider the LMM *P*_*τ*_ *y* = *Xβ* + *u* + *ϵ* with covariance matrix 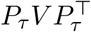. Plugging it in Equation 3, the log likelihood function changes to

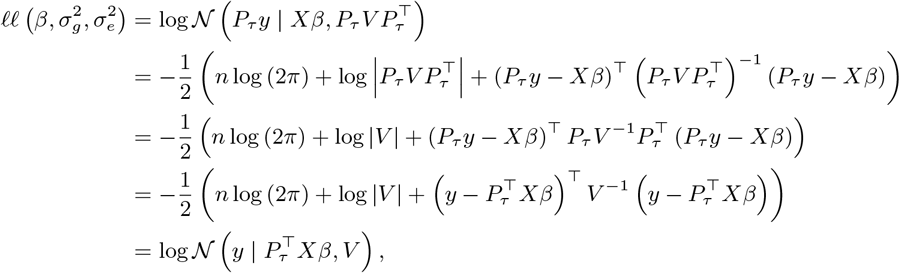

which is the log likelihood function of the LMM 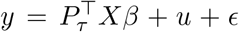 with covariance matrix *V*. Similarly, the restricted log likelihood changes to

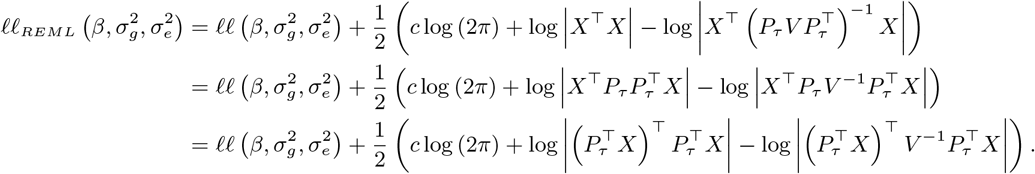

### 2.3 permGWAS2 architecture

Both of the above permutation strategies are computationally expensive and thus mostly infeasible in practice. To accelerate permutation-based approaches while efficiently computing univariate statistical tests, John *et al*. (2022) introduced batch-wise LMMs. Although the originally introduced `permGWAS` outperforms commonly used LMMs such as `EMMAX` (Kang *et al*., 2010) and `FaST-LMM` (Lippert *et al*., 2011) in terms of computational and statistical power, it still performs many redundant computations. In addition, batch-wise computations lead to an increased memory footprint compared to computing single univariate tests one at a time.

In the following, we first review the mathematical framework for batch-wise LMMs based on John *et al*. (2022), and then show how an elegant block matrix decomposition can be used to compute LMMs more efficiently.

#### Batch-wise linear mixed models

Let *b* be the number of genetic markers to test in parallel. Let *Z* ∈ ℝ^*n×*(*c*−1)^ be the matrix containing a column of ones for the intercept and all covariates, and let *s*_*i*_ ∈ ℝ^*n*^ be the *i*-th SNP for *i* ∈ {1, …, *b*}. Let *X*_*i*_ := [*Z, s*_*i*_] ∈ ℝ^*n×c*^ be the matrix of fixed effects consisting of *Z* and the SNP *s*_*i*_.

Let *E* := *D* + *δI*. Given the spectral decomposition *H* = *U* (*D* + *δI*) *U*^⊤^ = *UEU*^⊤^, let 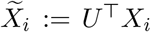 and 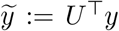. Denote by 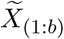 the 3-dimensional tensor in ℝ^*b×n×c*^ containing the matrices 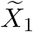to 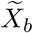 (see Figure 1 (A)). Further, for each matrix *A* ∈ ℝ^*n×d*^ denote by *A*_(*b*)_ the 3-dimensional tensor in ℝ^*b×n×d*^ obtained by stacking *b* copies of *A*. Then the effect sizes and the residual sums of squares for all SNPs can be computed simultaneously via

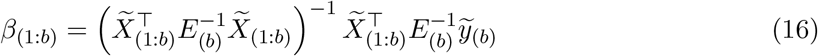

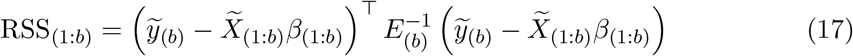

Now let *q* be the number of permutations. First, consider the original `permGWAS` permutation strategy (John *et al*., 2022), where we only permute the phenotype. Here, for each permutation ^*k*^*y* of *y* with *k* ∈ {1, …, *q*} let ^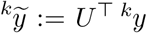^ and let ^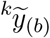^ denote the 3-dimensional tensor with *b* copies of ^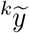^. By stacking ^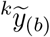^ for all *k* ∈ {1, …, *q*}, we get a 4-dimensional tensor 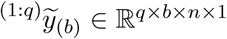. Similarly denote by ^(1:*q*)^*E*_(*b*)_ the 4-dimensional tensor in ℝ^*q×b×n×n*^ containing the 3-dimensional tensors ^*k*^*E*_(*b*)_ = (*D* + ^*k*^*δI*) _(*b*)_ for all *k* ∈ {1, …, *q*}. Further, for each 3-dimensional tensor *A* ∈ ℝ^*b×n×d*^ denote by ^(*q*)^*A* the 4-dimensional tensor in ℝ^*q×b×n×d*^ obtained by stacking *q* copies of *A*. Then the effect sizes and the residual sums of squares for *b* SNPs and *q* permutations can be computed simultaneously via

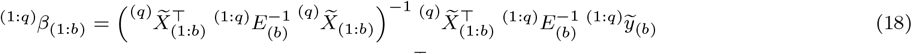

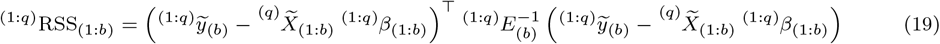

For the second permutation strategy, where both the phenotype vector and the covariance matrix are permuted, ^(1:*q*)^*β*_(1:*b*)_ and ^(1:*q*)^RSS_(1:*b*)_ can be computed similarly to the equations 18 and 19, except that instead of ^*k*^*y* we compute permutations ^*k*^*X*_*i*_ of *X*_*i*_:

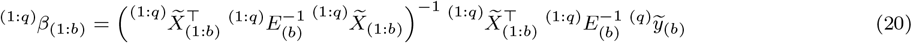

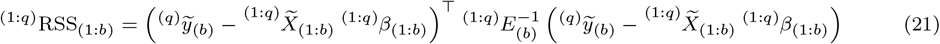

**Figure 1:**
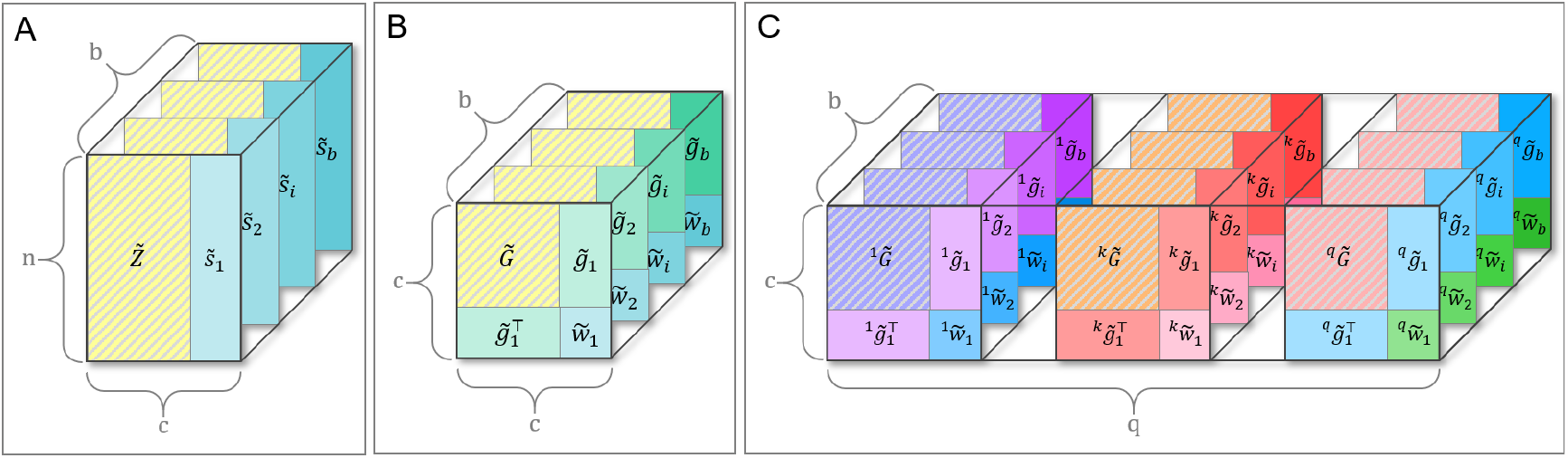
Schematic visualization of tensors and block matrices of the permGWAS2 architecture. Note that shaded areas are the same in each layer within a 3D tensor. (A) 3D tensor 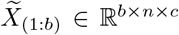 containing the fixed effects matrices 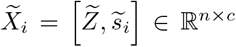 with SNPs *s*_*i*_ for *i* ∈ {1, …, *b*}. (B) 3D representation of block matrix structure with 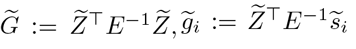 and 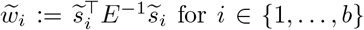 for *i* ∈ {1, …, *b*}. (C) 4D representation of permutation-based block matrix structure with 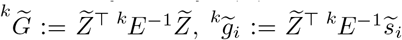 and 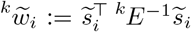 for *i* ∈ {1, …, *b*} an *k* ∈ {1, …, *q*}.

#### Efficient block matrix decomposition

In the following, we introduce a more efficient way to compute batch-wise LMMs and permutations, using an elegant block matrix decomposition that is applicable to both permutation strategies:

##### Lemma 2.

*Let Z* ∈ ℝ^*n×*(*c*−1)^, *s* ∈ ℝ^*n*^, *and let A* ∈ ℝ^*n×n*^ *be a symmetric matrix. Denote by X* := [*Z, s*] ∈ ℝ^*n×c*^ *the matrix obtained by concatenating the vector s to the matrix Z. Then the matrix X*^⊤^*AX has the following block structure*

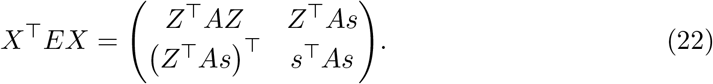

*Proof*. Let 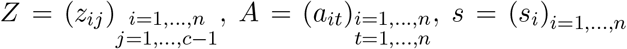 and 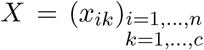.

Then

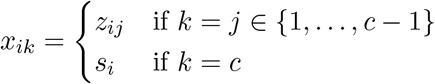

Denote the transposed matrices by 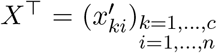 and 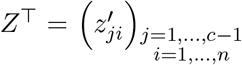, where 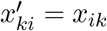 and 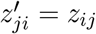. It follows that

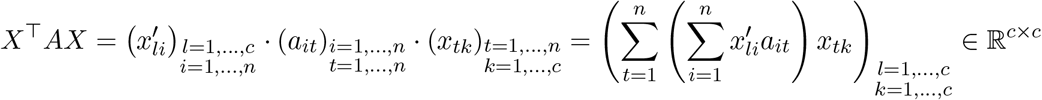

with

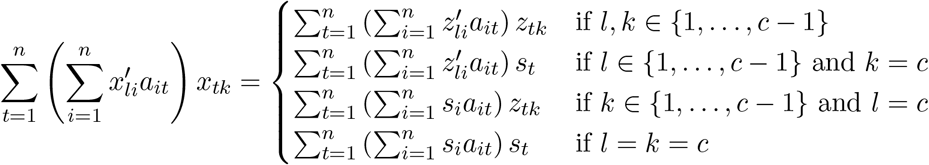

Since 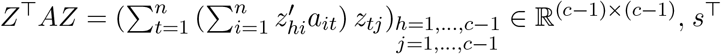 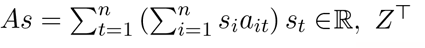 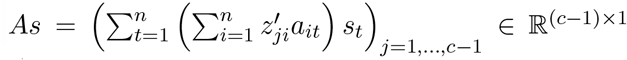 and *s*^⊤^*AZ* = (*Z*^⊤^*As)*^⊤^ ∈ R^1*×*(*c*−1)^, the claim follows.

Now le t 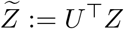 and 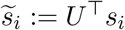 for all *i* ∈ {1, …, *b*}. Note that 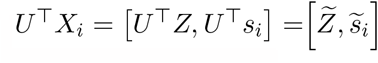. Denote by 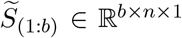 the 3-dimensional tensor containing the batch of *b* SNPs. Then by Lemma 2, we can decompose 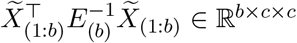 into the following block structure

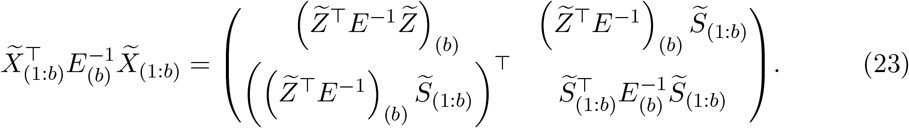

Since 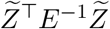 is independent of th e SNPs, we don’t need to recompute it for each SNP. Hence, we only need to compute 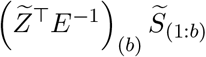 and 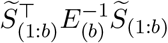 in batches and can assemble the block matrix afterward (see Figure 1 (B)). Similarly, one can show that

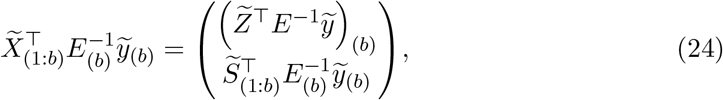

where 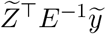 is independent of the SNPs. Again, only 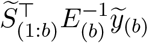 needs to be computed in batches.

Now when performing GWAS with permutations, we can decompose 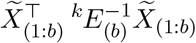 and ^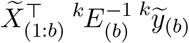^as in Equations 23 and 24 for each *k*. Since 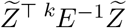 and 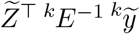*y* only depend on the permutation and not on the SNPs, we only need to compute them once per permutation (see Figure 1 (C)).

We call the new `permGWAS` version using this block matrix decomposition `permGWAS2`. Per default, `permGWAS2` uses the permutation strategy, where we permute both the phenotype and the covariance matrix. In the following, we refer to `permGWAS2` with the old permutation strategy, where we permute only the phenotype vector, as `permGWAS2(y)`.

### 2.4 Implementation & Availability

`permGWAS2` is implemented in Python3 as a standalone command-line tool. It uses `PyTorch` in addition to common scientific computing libraries such as `numpy`, `pandas`, and `scipy` to support multi-core and GPU usage as well as efficient tensor arithmetic. We support several common genotype and phenotype file formats, including PLINK, CSV, and HDF5. For the genetic similarity matrix, the user can either provide a pre-computed matrix or use the implemented realized relationship kernel, which is computed by `permGWAS2` by default. In addition, the user can specify covariates to account for certain fixed effects. To estimate the variance components, `permGWAS2` includes a custom optimization function using Brent’s method. For permutations, the user can choose between two strategies, the default strategy of permuting the phenotype vector and the covariance matrix (called `permGWAS2`), and the old strategy of only permuting the phenotype *y* (called `permGWAS2(y)`). To simplify the workflow, `permGWAS2` supports the use of YAML configuration files. Our framework also includes functions to visualize p-values as Manhattan or QQ-plots. It is also possible to include models other than LMMs within the `permGWAS2` framework. Our code is open source and publicly available on GitHub: https://github.com/grimmlab/permGWAS.

## 3 Results & Discussion

To compare the performance of `permGWAS2` with block matrix decomposition with the original `permGWAS` version (John *et al*., 2022), we performed several runtime experiments. Additionally, we compared the runtime of both versions with the two commonly used LMMs `EMMAX` (Kang *et al*., 2010) and `FaST-LMM`. (Lippert *et al*., 2011), for which we used the binary C/C++ implementations. The runtime comparisons were performed on a machine running Ubuntu 22.04.2 LTS with a total of 52 CPUs, 756 GB of memory, and 4 NVIDIA GeForce RTX 3090 GPUs with 24 GB of memory each. For our experiments, we used dedicated Docker containers where we limited the number of CPUs to one core and used a single GPU. We analyzed the runtime in terms of the number of markers, the number of samples, and the number of permutations. To do this, we generated synthetic data with varying numbers of samples and markers by up- and down-sampling a flowering time related phenotype from *Arabidopsis thaliana* and the corresponding genotype matrix (The 1001 Genomes Consortium, 2016). For each experiment, we took the average of three runs.

To evaluate the effect of increasing the number of markers, we first fixed the number of samples at 1000 and varied the number of SNPs between 10^4^ and 5 *×* 10^6^. As shown in Figure 2 (A), both `permGWAS` and `permGWAS2` show similar performance with about 14 min for `permGWAS2` and 17 min for `permGWAS` for 5 million SNPs, visibly outperforming `EMMAX` and `FaST-LMM`. In the second experiment, we fixed the number of markers at 10^6^ and varied the number of individuals between 100 and 10^4^ to analyze how the runtime depends on the number of samples. Again, `permGWAS2` slightly outperforms `permGWAS` with about 55 min compared to 67 min for 10k samples, both at least an order of magnitude faster than the comparison partners as summarized in Figure 2 (B).

**Figure 2:**
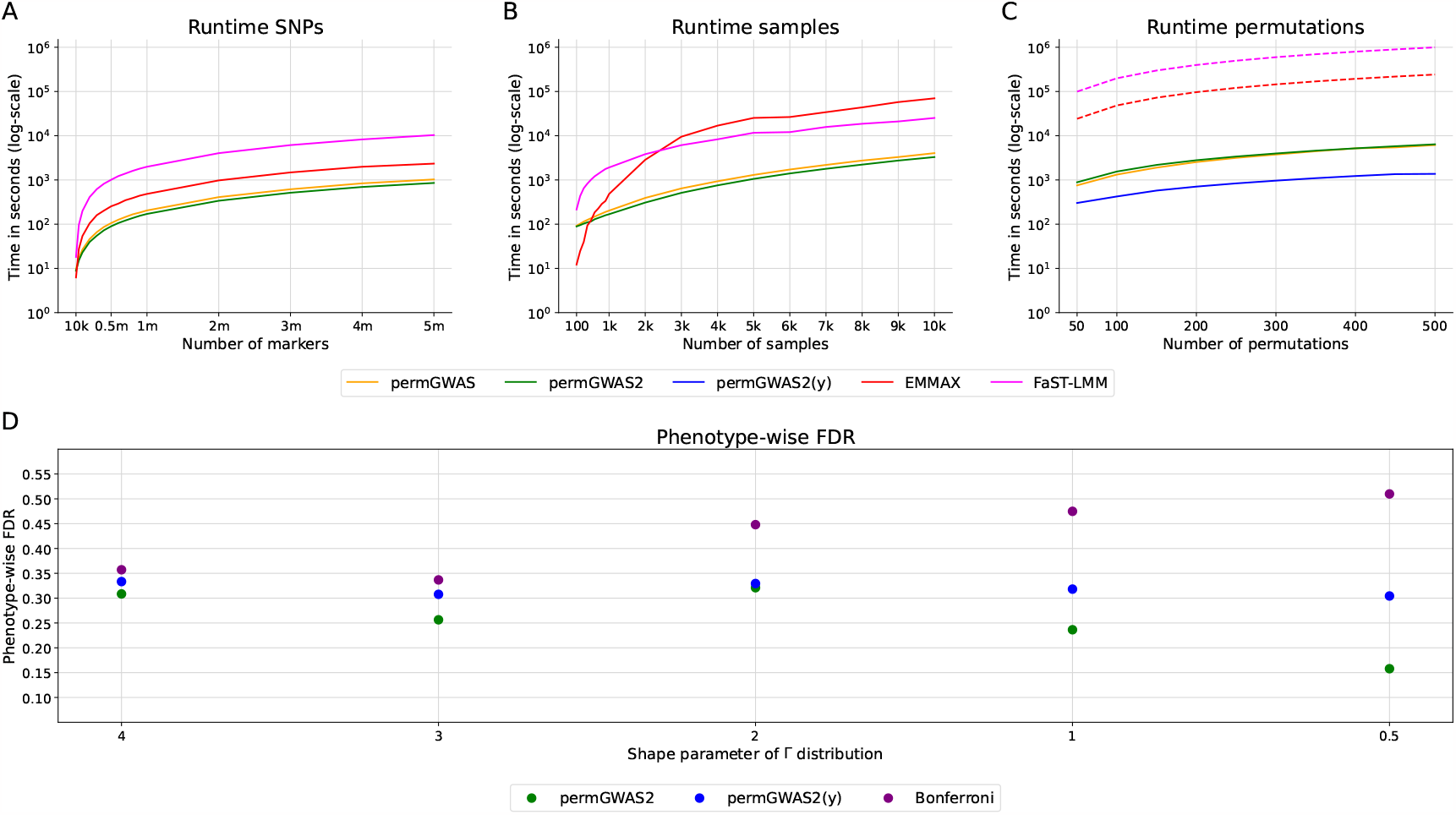
Performance analysis of permGWAS2. (A) - (C) Runtime comparisons of `permGWAS2` vs. `permGWAS`, `EMMAX` and `FaST-LMM`. Note all axes are log-scaled. (A) Computational time as function of number of SNPs with fixed number of 1 000 samples. (B) Computational time as function of number of samples with 10^6^ marker each. (C) Computational time as function of number of permutations with 1 000 samples and 10^6^ marker each. Dashed lines for `EMMAX` and `FaST-LMM` are estimated based on the computational time for 1 000 samples and 10^6^ markers times the number of permutations.`permGWAS` and `permGWAS2` were run on a single GPU. `permGWAS2(y)` refers to `permGWAS2` with the same permutation strategy as in the previous `permGWAS` version where only the phenotype vector is permuted. (D) Phenotype-wise FDR for Bonferroni and permutation-based significance threshold with `permGWAS2` and `permGWAS2(y)`.

Finally, we again set the number of samples to 1000 and the number of markers to 10^6^, and ran between 10 and 500 permutations. We compared not only `permGWAS` with `permGWAS2`, but also with the old permutation strategy `permGWAS2(y)`, which only permutes the phenotype vector. Since `EMMAX` and `FaST-LMM` do not support permutations, we took the runtime from the previous experiments and approximated the permutation-based time by multiplying it by the number of permutations. However, this is only an estimate of the runtime without taking into account the time needed for additional pre- and post-processing steps to prepare the permutations. For 500 permutations the performance of `permGWAS2` is similar to that of `permGWAS` with 107 min, and 102 min, respectively (see Figure 2 (C)). This is remarkable considering that `permGWAS2` permutes the fixed effects matrix for all markers instead of only permuting the phenotype vector. If we use this simpler permutation strategy with `permGWAS2(y)`, 500 permutations take around 23 min, so less than a quarter than the original `permGWAS`. In comparison, the approximated runtimes of `EMMAX` and `FaST-LMM` are close to 3 days and more than 11 days, respectively.

To compare the two permutation methods `permGWAS2` and `permGWAS2(y)`, we simulated artificial phenotypes for 200 random *A. thaliana* individuals based on fully imputed genotype data with approximately 2.8 million SNPs (Arouisse *et al*., 2020). We first computed the realized relationship kernel as a kinship matrix and its Cholesky decomposition *K* = *CC*^⊤^. To simulate the polygenic background we chose a random vector *u* ∈ R^*n*^ where each element was drawn from a Gaussian distribution with zero mean and a variance of 1 and multiplied it with *C* such that Var(*Cu*) = *K*. Next, we added random noise drawn from a gamma distribution such that the noise contributed 70% to the total phenotypic variance. Finally, we added a causal SNP with a minor allele frequency greater than 5% and an effect size that explained about 20% of the phenotypic variance. To simulate differently skewed phenotypes, we used shape parameters of 0.5, 1, 2, 3 and 4 for the gamma distribution resulting in 5 different simulation settings. For each setting, we generated 100 simulations and ran `permGWAS2` and `permGWAS2(y)` with 500 permutations each. Since linkage disequilibrium decays on average within 10 kbp in *A. thaliana* (Kim *et al*., 2007), we defined a phenotype as a true positive (TP) if any marker within a 10 kbp window around the causal SNP was deemed significant. If a marker outside this window was significant, we classified the phenotype as a false positive (FP). Thus, a phenotype could be both TP and FP at the same time. We then calculated for each of the five simulation settings the phenotype-wise false discovery rate 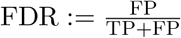 for both permutation-based and Bonferroni threshold.

Our experiments show that the phenotype-wise FDR for `permGWAS2(y)` is more or less stable over all simulation settings (see Figure 2 (D)). In contrast, the FDR for `permGWAS2` decreases for smaller shape parameters, i.e., when the phenotypes become more skewed. Thus, by taking the population structure into account when permuting, we are able to better control the phenotype-wise FDR. Remarkably, the FDR with Bonferroni threshold increases up to 50% for the most skewed phenotypes, meaning that we find as many TP as FP associations.

## 4 Conclusions

We introduced `permGWAS2`, an improved and accelerated modification of `permGWAS` (John *et al*., 2022). By employing a block matrix decomposition, `permGWAS2` optimizes computations and significantly reduces redundancy. In particular, our framework incorporates an improved permutation strategy that accounts for population structure during permutation. Our results demonstrate that `permGWAS2` provides a more streamlined approach with a lower false discovery rate compared to its predecessor and the traditional Bonferroni correction. This advancement represents a significant refinement in computational efficiency and statistical robustness for GWAS.

